# Transcriptomic signatures of host immune responses in aphthous ulcers, the earliest lesions of Crohn’s disease, suggest that a bacterial invasive challenge, rather than global dysbiosis, is the initiating factor

**DOI:** 10.1101/2024.09.28.615566

**Authors:** Phillip J Whiley, Ojas V A Dixit, Mukta Das Gupta, Hardip Patel, Guoyan Zhao, Susan J Connor, Kim M Summers, David A Hume, Paul Pavli, Claire L O’Brien

**Author notes:** **Correspondence** Phillip J Whiley. The Canberra Hospital, Yamba Drive, Garran, ACT, Australia 2605.

## Abstract

Crohn’s disease is a chronic, transmural inflammatory disease of the human gut. Changes in the faecal microbial composition and a reduction in species diversity (dysbiosis) are consistent features in studies of Crohn’s disease patients, but whether dysbiosis is a cause or consequence of inflammation remains unresolved. Genetic susceptibility also plays a role in the development of Crohn’s disease and has been linked to genes involved in recognition of intestinal bacteria by cells of the mononuclear phagocyte system. The earliest visible lesions in Crohn’s disease are aphthous ulcers, overlying Peyer’s patches and lymphoid follicles. To identify mechanisms underlying the earliest stages of disease initiation we compared gene expression in aphthous ulcers, Peyer’s patches, inflamed and endoscopically normal mucosa from patients and controls using total RNA-seq.

The resulting data was subjected to network analysis to identify co-regulated gene expression signatures of cell types and processes. These results were compared to recent single cell RNA-seq analysis of intestinal macrophages in normal and diseased mucosa. The analysis of aphthous ulcers revealed signatures of epithelial stress and antimicrobial defence, plasma cell activation and immunoglobulin production, monocyte recruitment, inflammatory gene expression and induction of interferon-γ and downstream target genes. These signatures were not present in the normal appearing mucosa adjacent to aphthous ulcers which were similar to healthy control mucosa.

We conclude that the initial lesion in Crohn’s disease arises from an invasive bacterial challenge leading to intense activation of multiple host defence pathways rather than the breakdown of epithelial barrier integrity and widespread bacterial translocation.

## 1. Introduction

Crohn’s disease (CD) is a debilitating chronic inflammatory condition affecting the gastrointestinal tract. Changes in the microbial composition and a reduction in species diversity (dysbiosis) are consistent features in studies of Crohn’s disease patients. It is postulated that these changes affect epithelial barrier integrity permitting translocation of bacteria and their products, and/or compromise the host inflammatory response^1^.

Genetic susceptibility also plays a role in the development of Crohn’s disease: based upon results from genome-wide association studies (GWAS), it is widely believed that heritable disease susceptibility is linked to hyper-responsiveness of cells of the monocyte-macrophage lineage to gut microbiota^2^. Very early onset forms of IBD inherited in a Mendelian fashion are commonly due to mutations in monocyte-macrophage-expressed genes, and CD susceptibility loci identified by GWAS are strongly-enriched for regulatory elements associated with monocyte to macrophage differentiation and/or activation^3^.

In an effort to identify inflammatory mechanisms and therapeutic targets, the past 10 years have seen numerous published transcriptomic analyses using either total RNA sequencing (RNA-seq) or single cell RNA sequencing (scRNA-seq), comparing affected and unaffected areas of patient GI tract^3-8^. Not surprisingly, many immune response-related genes are overexpressed in inflamed bowel.

The potential weakness of these studies is the focus on established lesions in inflamed mucosa. A seminal study of post-operative recurrence in Crohn’s disease patients demonstrated that aphthous ulcers (AU) are the earliest detectable lesions^9^. These lesions overlie the follicle associated epithelium (FAE) of the small bowel (Peyer’s patches, PP) and large bowel (lymphoid follicles) and were detected in 70% of patients with Crohn’s disease ^9^. There is a predictable sequence from AU to typical ileitis ^10^. Radiological and endoscopic studies^11^ showed that around 30% of patients with an “incidental” finding of isolated terminal ileal aphthous ulceration subsequently develop Crohn’s disease.

PP and colonic lymphoid follicles are secondary lymphoid structures that facilitate the interaction between gut antigens and antigen-specific lymphocytes leading to the induction of intestinal immunoglobulin and other responses ^12^. Luminal antigens are delivered by specialized microfold (M) epithelial cells within follicle-associated epithelium to underlying antigen-presenting cells to initiate immune responses. M cells also provide a route of entry for various pathogens, such as *Yersinia pseudo-tuberculosis, Salmonella typhimurium* and *Shigella flexneri*^13^. Adherent-invasive *E. coli* adhere to follicle-associated epithelium and were more commonly isolated from mucosal biopsies of patients with Crohn’s disease than controls^14^. Increased uptake of non-pathogenic *E. coli* has also been demonstrated in longstanding Crohn’s disease but not ulcerative colitis ^15^. These studies led to speculation that the microbial triggers of Crohn’s disease gain access to the intestinal lamina propria via PP^16^.

In order to identify mechanisms underlying the earliest phases of CD, we focused on aphthous ulceration in a unique group of Crohn’s disease patients on no treatment. We compared gene expression and microbial profiles of AU and adjacent mucosa, and PP and adjacent mucosa from controls. To compare with established disease and to highlight responses that are specific to inflamed PP, we also profiled involved and adjacent mucosa and mesenteric lymph nodes from patients with active disease. In parallel, we characterised the mucosa-associated virome and microbiome of the earliest identifiable lesion in CD to identify candidate triggers that could be associated with initiation of disease. We conclude that AU are associated with evidence of microbial invasion, B cell activation and antibody production, an epithelial stress response and activation of recruited monocytes to produce inflammatory cytokines. These changes were not seen in the adjacent normal appearing mucosa. Network analysis of the data highlights informative co-expression signatures of different mucosal cell populations and is discussed in comparison to recent large scRNA-seq datasets from normal and inflamed human intestine ^17^.

## 2. Materials and Methods

### 2.1 Human subjects

The study was approved by the ACT Health Human Research Ethics Committee (ETH.5.07.464), and Australian National University Human Ethics Committee (2012/596) and written informed consent was obtained from all subjects. Crohn’s disease diagnoses were based on typical clinical presentation, and endoscopic and histological findings. Biopsies used for the analysis were obtained from the terminal ileum only, for both patients and controls, unless otherwise stated. Biopsies were also sent for histological assessment in parallel to confirm the endoscopic findings. Excluding patients who underwent surgical resection, patients were not on pharmacological treatment for CD. Details of the patients are provided in **Table S1**.

### 2.2 RNA extraction and NextSeq 500 sequencing

RNA was extracted from: 1) biopsies of AU and adjacent endoscopically unaffected mucosa in CD patients; and PPs and adjacent mucosa in controls; 2) biopsies of established inflammation comprising active ulceration and adjacent endoscopically unaffected mucosa in CD patients; and healthy mucosa from non-IBD subjects; 3) resected specimens of ulcerated mucosa, adjacent unaffected mucosa and lymph nodes from CD patients; and resected mucosa and lymph node specimens from non-IBD subjects.

RNA was extracted using Qiagen RNeasy Mini kits (Qiagen). Each biopsy was mechanically homogenized using a Qiagen TissueLyser II, and an on-column DNase digestion step was performed as per the kit protocol. An aliquot of denatured RNA was quantified using an Agilent 2100 Bioanalyzer with RNA 6000 Nano LabChips (v2). NextSeq 500 libraries were prepared at the Biological Research Facility (Australian National University) using 1 ug RNA for each sample. Truseq stranded RNA LT kits, and NextSeq 500/550 High output kits (Illumina), in a 150 × 150 bp (paired-end) format were used as per the manufacturer’s instructions. Whole genome transcripts were assessed for quality using FASTQC, trimmed using Trimmomatic, and aligned to the human reference genome (Hg38) using subread mapper. Fragment counts were obtained using featureCount, and expression values normalised using the trimmed mean of M-values normalisation method (TMM)^18^. Reads that did not map to the human genome in RNA-seq data from each sampled tissue were used to mine virus sequences using the VirusSeeker pipeline^19^.

For qRT-PCR validation of gene expression differences, RNA was extracted from RNAlater®-stabilised frozen biopsies or approximately 30 mg of resected bowel tissue, using RNeasy Mini Kits (Qiagen). Samples were treated for DNA contamination using RNase-Free DNase (Qiagen). cDNA was synthesised using 500 ng of total RNA and Superscript III cDNA First Strand Synthesis kit (Invitrogen) with random primers. Quantitative PCR was carried out using a Quantstudio 12K system, SYBR Green (Invitrogen). The primers listed in **Table S2**. PCR reactions were conducted under the following conditions: 95°C for 3 min, followed by 45 cycles of 95°C for 10 sec, 60°C for 20 sec and 72°C for 20 sec, one cycle of 95°C for 1 min and 60°C for 1 min, and 71 cycles from 60°C to 95°C for melt curve analysis with 0.5°C increases. Two experiments were run in duplicate and analysed using *GAPDH* and *ACTB* as reference genes. Taqman™ fast advanced protocol (Applied Biosystems) was used for *NOD2* (Hs01550753_m1) and *LILRA2* (Hs01597933_g1). All experiments were carried out using 2 biological and 3 experimental replicates. All authors had access to the all data and have reviewed and approved the final manuscript.

### 2.3 Data analysis

#### 2.3.1 Transcriptional network analysis

Transcripts that were not detected in at least 6 samples were removed to minimise stochastic sampling errors. Network analysis was performed using the program, BioLayout (http://biolayout.org) or a further development of this platform Graphia (https://graphia.app/) ^20^. By contrast to the widely-used WGCNA approach ^21^ this network method generates an all vs. all correlation matrix to which it applies a correlation threshold cut-off removing outliers. This thresholded matrix is used to generate a true correlation graph from which co-expression modules are then defined using the Markov clustering (MCL) algorithm, which further refines and removes outliers. The outcomes are similar to WGCNA, but modules defined by this approach are biologically enriched over those defined by WGCNA ^20^. Pairwise Pearson correlations (*r*) were calculated between samples to produce a sample-to-sample correlation matrix and inversely between all pairs of genes to produce a gene-to-gene correlation matrix. Gene co-expression networks were generated from the matrix, in which nodes represent genes and edges represent correlations between nodes above a defined correlation threshold. An *r* value threshold of 0.75 was used for all gene-to-gene analyses unless otherwise stated. Further analysis used the Markov algorithm (MCL) with an inflation value of 2.0 to identify groups of highly connected genes within the overall topology of the network.

Differential gene expression analyses were also performed using generalized linear models in edgeR^18^, with a false discovery rate (FDR) applied after correcting for multiple hypothesis testing using the Benjamini and Hochberg method^22^.

Gene ontology (GO) terms were derived from the Gene Ontology Resource (http://geneontology.org) using the PANTHER overrepresentation test (PANTHER).

#### 2.3.2 Microbiome analysis

The V3-V4 region of the 16S rRNA gene was amplified from RNA derived from tissue biopsies and the corresponding amplicon library was sequenced on an Illumina MiSeq platform. Qiime2-2019.10 was used for the generation of alpha and beta diversity metrics, distance analyses, and for visualising the data.

## 3. Results

### 3.1 Network analysis

Total RNA-seq data was generated from a total of 48 biopsies, including AU; control PP; normal-appearing mucosa from CD patients and healthy controls; inflamed and uninvolved mucosa and lymph nodes from CD patients with active disease. On average we obtained 36 million reads per sample of which 87% were retained for downstream processing, and 93% of these mapped to the human genome. Gene expression was quantified as described in Materials and Methods. The complete set of quantified expression data is provided in **Table S3**. Aside from the differences in location (PP, mucosa, lymph node) individual biopsies from the same location also differ in cellular composition and activation status. Network analysis exploits this diversity to enable extraction of sets of correlated transcripts that define cell types, sub-tissue locations and/or cellular processes. This approach has been used previously to identify cellular signatures in very large datasets including those derived from human tumours ^23^.

A gene-centred network (GCN) was generated at a threshold Pearson correlation of r=0.75 and an MCL inflation value (defining granularity) of 2.0. As in previous studies, the threshold was chosen empirically to maximise the number of nodes and minimise the number of edges. **Table S4** contains the lists of genes and the average expression profiles of clusters of interest and annotation of the clusters highlighting the power of the analysis to extract informative co-expression signatures of different cell types and processes. Amongst the clusters, a subset showed a pattern of shared expression corresponding to inflammatory state. **Clusters 4, 8, 9, 14** and **27** distinguished AU from PP; of these, gene expression in **Cluster 8** and **14** was also increased in inflamed mucosa and lymph nodes from CD patients (**Figure 1**). These 6 Clusters were assessed for enrichment with known and candidate susceptibility genes ^24^. Taken in combination, **Clusters 4, 8, 9, 14, 16** and **27** contained 30 of 231 known IBD susceptibility genes and 28 of 153 monocyte candidate genes respectively. The smaller **Cluster 37** contains KIT and several mast cell-specific transcripts but was not associated with disease state. To determine whether or not there was epithelial cell injury in the mucosa adjacent to AUs, we examined the expression of 23 epithelial-cell specific genes (Table 1) (bioGPS.org), and compared AUs, the corresponding adjacent normal appearing mucosa, PPs and the corresponding adjacent mucosa (Figure 2). There was no evidence for epithelial injury in the mucosa adjacent to PPs or AUs to suggest a more global microbial insult.

**Table 1.**
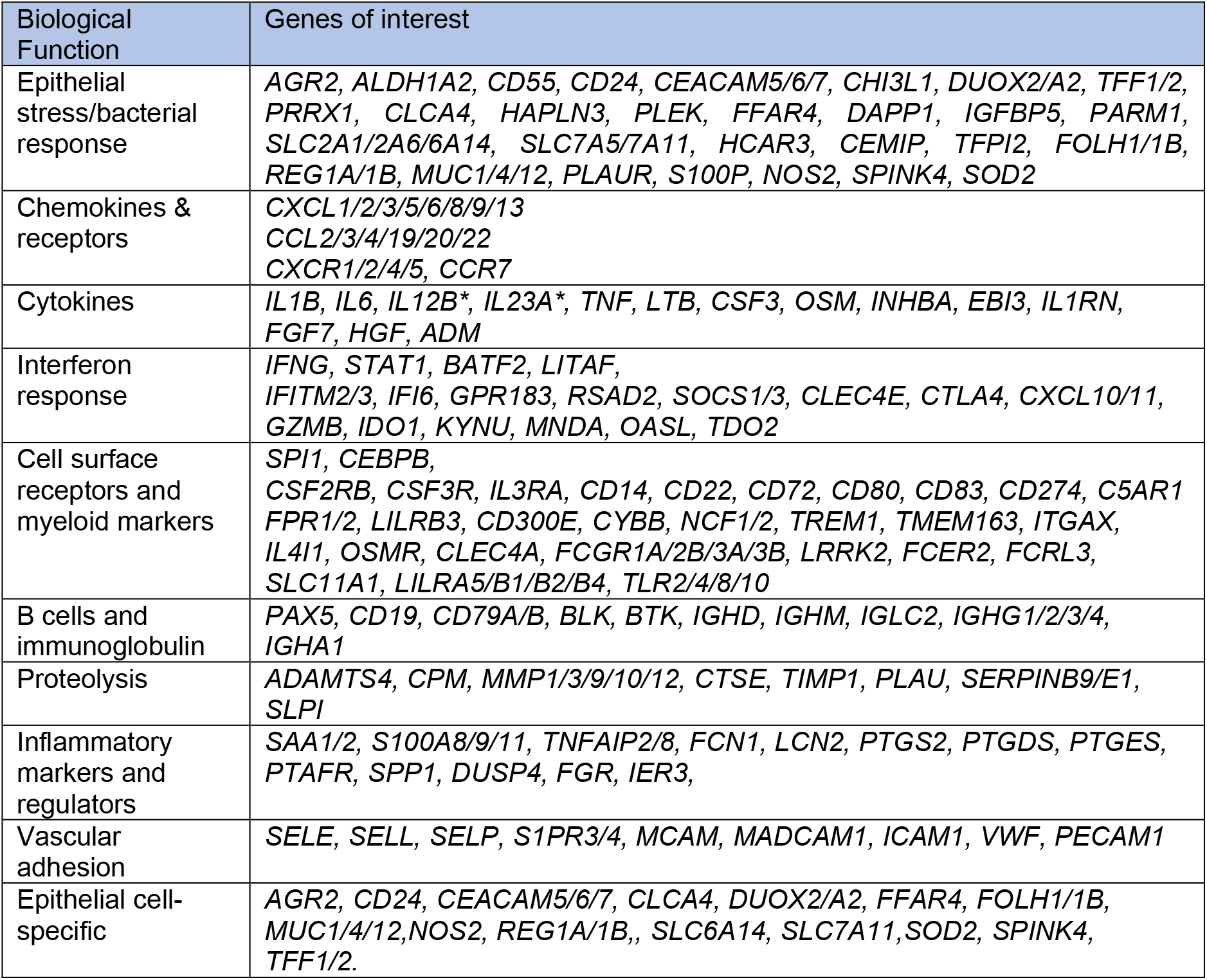
Functional classification of transcripts up-regulated in aphthous ulcers compared to Peyer’s patches. (Full list of regulated transcripts is shown in Table S4).

**Figure 1:**
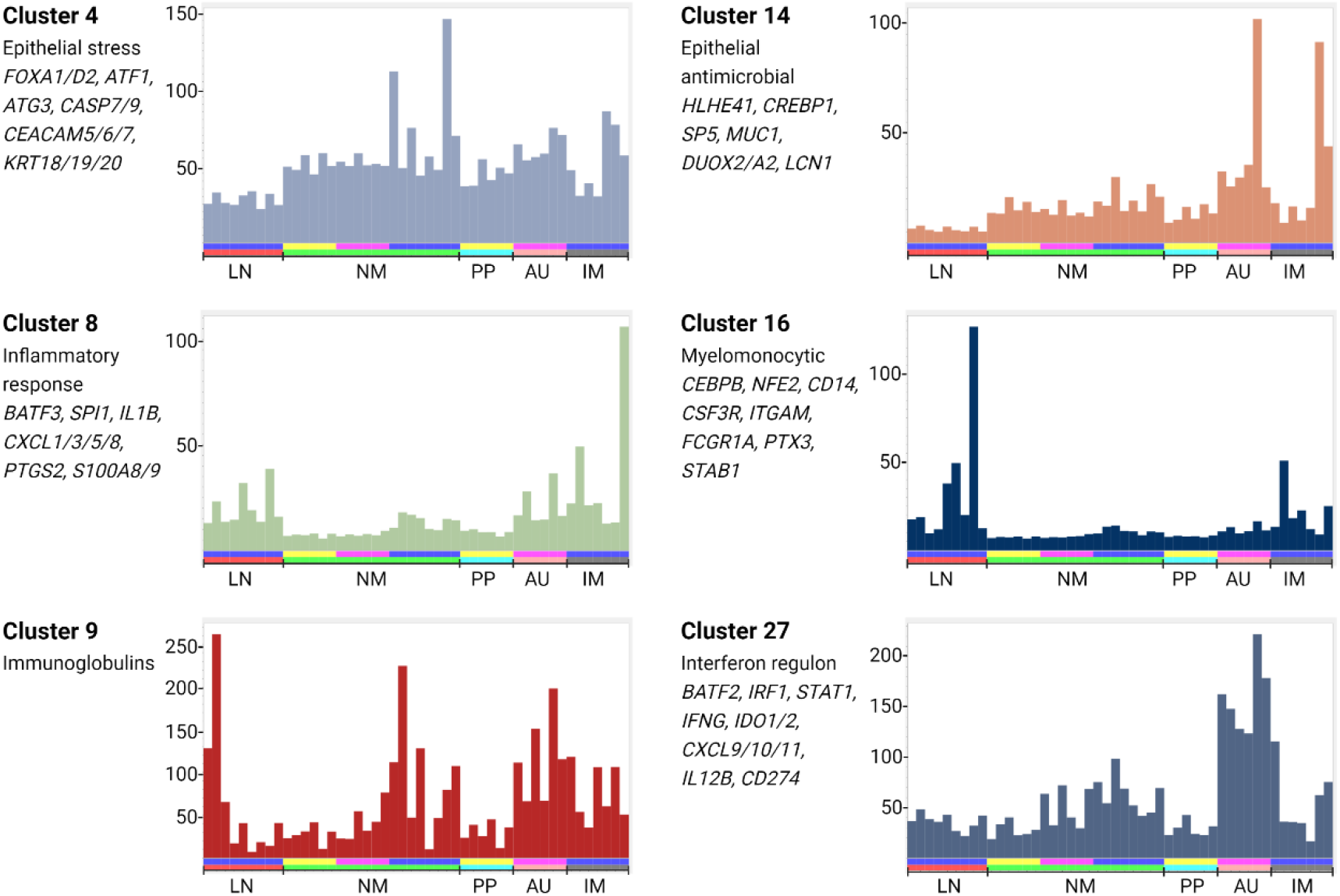
Co-expression clusters identified from gene-centred network analysis. Total RNA-seq data was generated from 48 biopsies, including aphthous ulcers (AU); control Peyer’s patched (PP); normal-appearing mucosa from CD patients and healthy controls (NM); inflamed mucosa (IM) and lymph nodes (LN) from CD patients with active disease. A gene-centred network was generated using Biolayout as described in Materials and Methods. The full list of clusters is provided in **Table S4**. Histograms in this figure show the average expression profile of the set of transcripts in clusters of specific interest, an indication of the shared pattern of expression. Y axis: average expression (TPM). X axis : samples. Upper bar: blue - CD patients with active disease; pink - aphthous ulcer CD patients; yellow - controls. Lower bar: red – lymph node; green – normal mucosa; turquoise – Peyer’s patch; salmon – aphthous ulcer; grey – involved mucosa.

**Figure 2:**
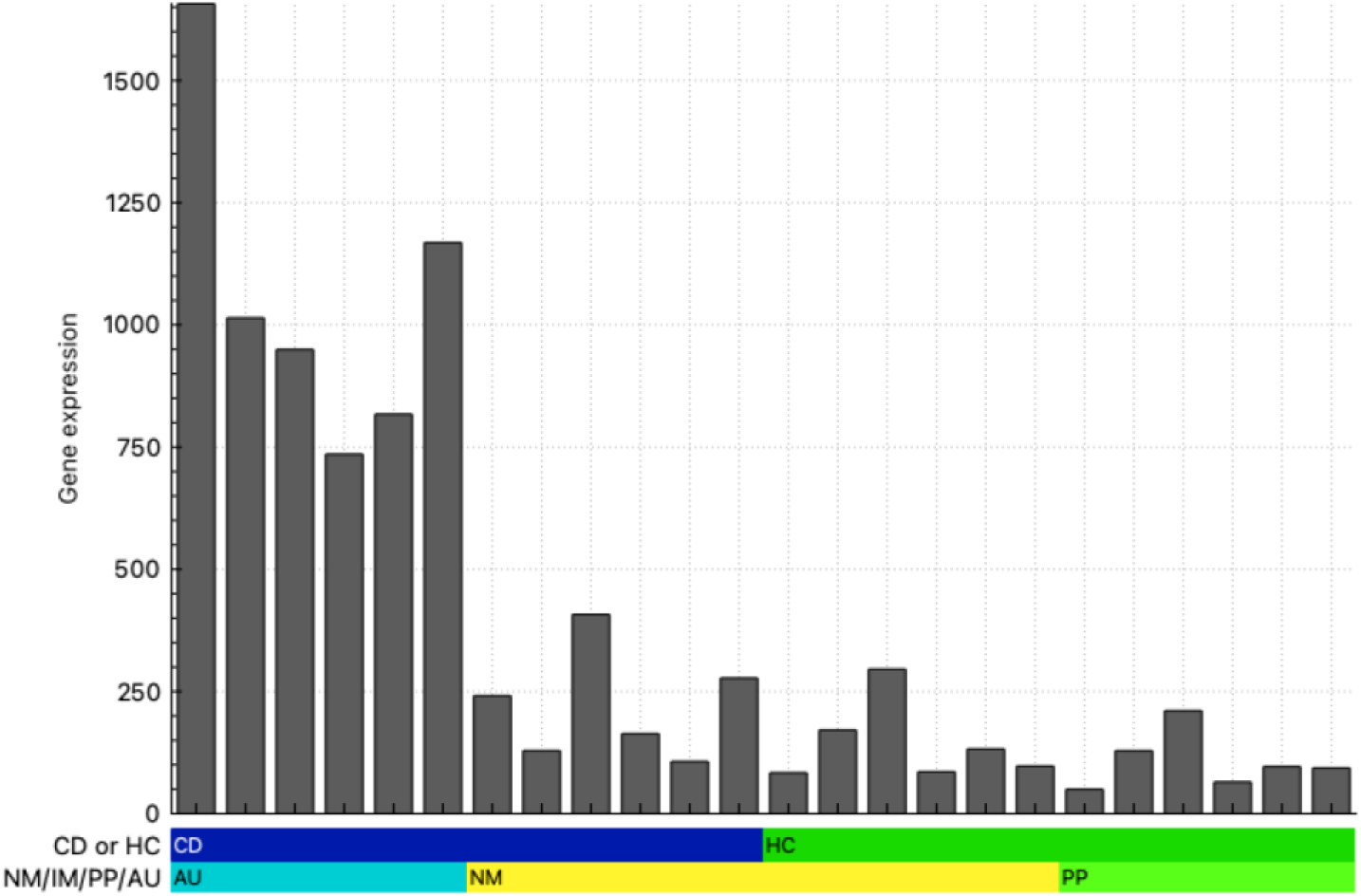
Epithelial cell-specific gene network analysis. Total RNA-seq data was generated from 24 biopsies, including aphthous ulcers (AU); Peyer’s patches (PP); and normal-appearing mucosa from CD patients and healthy controls (NM). A gene-centred network was generated using Biolayout as described in Materials and Methods. Histograms show the average expression profiles of the set of gene transcripts of interest (Table 1). Y axis: average expression (TPM). X axis: samples. Upper bar: blue - CD patients; green - healthy controls. Lower bar: turquoise – AU samples from six CD patients; yellow – normal-appearing mucosa from the six corresponding CD patients in sequence and normal mucosa from six healthy controls; lime green - Peyer’s patches from the corresponding six healthy controls in sequence.

### 3.2 Differential gene expression

Network analysis groups transcripts that share a pattern of expression, but the basal level of expression and fold differences vary for transcripts within a cluster. Furthermore, depending upon the correlation threshold, some additional differentially-regulated genes will form smaller clusters based upon idiosyncratic differences between individual biopsies. **Table S5A** shows the results of a conventional differential gene expression analysis, comparing AU and PP. Excluding transcripts that were not detected at >5TPM on average in AU biopsies, there were 494 transcripts increased significantly in AU versus PP. These are broadly annotated in a ranked list in **Table S5B. Table 1** shows a curated set of genes that were up-regulated in AU excluding Ig variable genes and curated into broad functional classes based upon the known gene function. IL23 (subunits encoded by *IL12B* and *IL23A*), which is strongly implicated in IBD pathology ^25^, fell below the 5TPM detection threshold, but both transcripts were increased >5-fold in AU versus PP.

To validate the patterns of altered gene expression seen in the RNA-seq analysis, we performed qRT-PCR analysis on 13 transcripts selected from **Table 1**, comparing AU and adjacent mucosa in terminal ileum and colon obtained from a separate cohort of CD patients. Transcripts were chosen to sample novel genes of interest across a range of biological functions and distinct clusters, including myeloid markers (*TREM1, FPR1, OSMR, LILRA2, NOD2*), inflammatory diagnostic markers (*S100A12, FCN1*), antibody (*IGHA1*), cytokine (*OSM*), antigen processing (*TAP1*) and epithelial response (*CEACAM4/6*). *TGM2* was recently identified as a novel marker of epithelial responses in experimental and human colitis ^26^. All 13 transcripts were significantly up-regulated in AU in the terminal ileum. The pattern was more variable in the colon, where *IGHA* was not detected and *LILRA2* and *NOD2* were unaffected (**Figure 3**). To extend the analysis we compared the expression of the same set of transcripts in actively inflamed tissue to either uninflamed mucosa adjacent to AU or mucosa from non-IBD controls in terminal ileum (**Figure S1**) or in colon (**Figure S2**). All transcripts except *TAP1* distinguished active inflammatory mucosa from inactive and/or control mucosa in terminal ileum. However, in the colon the association with involved mucosa was less evident. Whereas 9/13 transcripts (excluding *TAP1, TGM2, IGHA1* and *CEACAM6*) were up-regulated in involved compared to uninvolved mucosa, *FPR1, OSMR* and *S100A12* were also expressed in normal mucosa, albeit of unrelated individuals.

**Figure 3.**
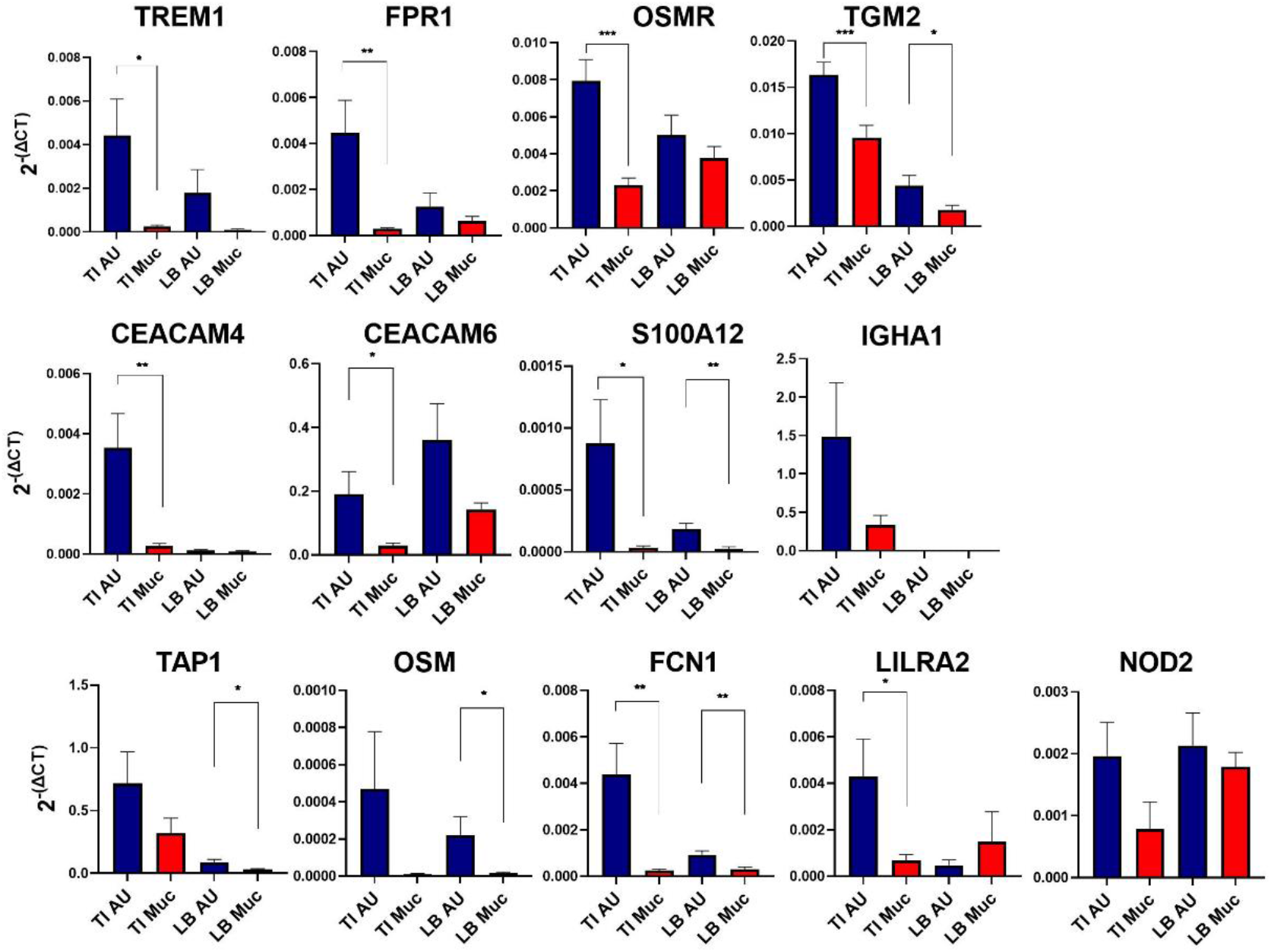
Expression of selected transcripts in aphthous ulcers and adjacent mucosa. mRNA was isolated from terminal ileum aphthous ulcers (TI AU), adjacent terminal ileum normal appearing mucosa (TI Muc), large bowel aphthous ulcers (LB AU) and large bowel normal appearing mucosa (LB Muc) and expression of the indicated transcripts was determined by qRT-PCR as described in Materials and Methods. ^*^P<0.05, ^**^P<0.01, ^***^P<0.001 for pairwise comparison of affected versus unaffected tissue.

### 3.3 Microbiome of aphthous ulcers and Peyer’s patches

Ryan *et al*. ^27^ reported that mucosa of patients with IBD trended towards lower microbiota diversity compared to healthy controls, but no individual taxa distinguished inflamed from non-inflamed sites within the same individual. The profound induction of proinflammatory cytokines and chemokines (**Cluster 8**) and genes associated with bacterial invasion (**Cluster 4**) in the AU compared to PP strongly suggests a response to a pathogen. We used VirusSeeker to examine the 7% of transcript reads that did not map to the human genome. There was no virus in common to all AU or lymph nodes from patients with Crohn’s disease. We detected human herpes virus 4 in one CD AU, one CD lymph node, and the mucosa of three CD patients, as well as one control lymph node (data not shown).

Naftali *et al*. ^28^ suggested that *Faecalibacteria* were strongly reduced in the microbiome of patients with ileal CD compared with CD with colonic involvement. The V3-V4 region of the 16S rRNA gene was amplified from RNA of the six AU that underwent RNA-seq, an additional six AU from different patients, and five of the six PP. A similarity of percentages analysis of the 10 most abundant genera in the AU and PP microbiomes is provided in **Table S6**. Notwithstanding the small group size, consistent with previous literature on the gut microbiome in CD^16, 29^ we observed a small depletion in *Faecalibacterium*. In contrast to previous literature, we saw a small depletion of *Escherichia_Shigella* in AU when compared to PP. As a group, the microbiomes of AU were less diverse than those of PP, when calculated using various indices (Chao1, p=0.03; Shannon, p=0.03; and, Fisher, p=0.03).

**Figure 4.**
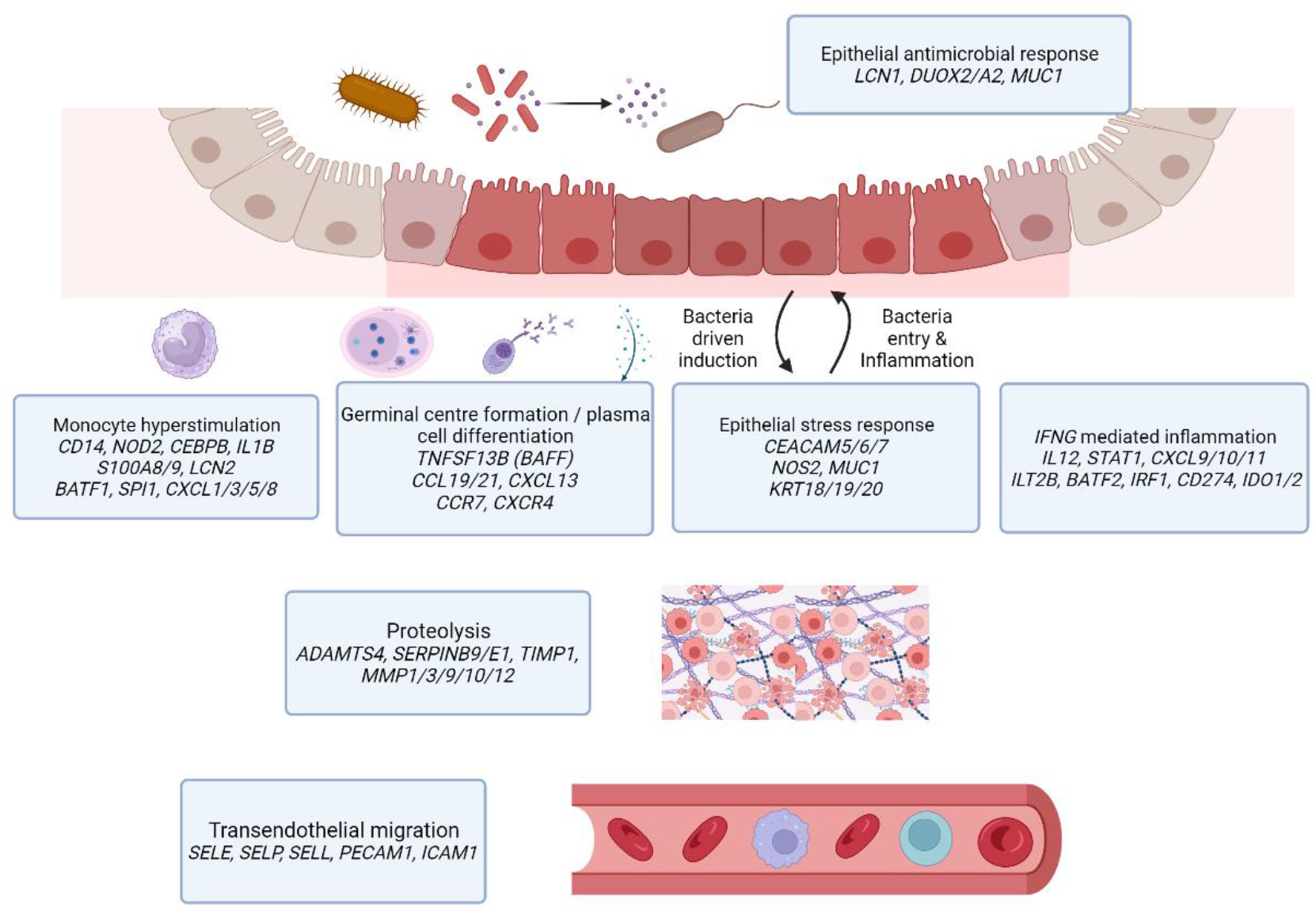
Inflammatory response pathways informed by RNAseq cluster analysis of aphthous ulcers early in Crohn’s disease inflammation. At the mucosa and lamina propria, highly expressed genes formed distinct functional groups suggestive of bacterial antigen induced epithelial stress, monocyte hyperstimulation, IFNG priming and plasma cell differentiation along with germinal centre formation. Distinctly up-regulated subsets of genes were observed for endothelial cell migration and proteolysis functions.

## 4. Discussion

IBD is a chronic, debilitating condition that tends to manifest in early adulthood and runs a relapsing and remitting course. Despite a diverse range of monoclonal antibodies targeting specific inflammatory pathways, resistance to therapy is an ongoing clinical problem^30^. The need for identification of new targets for treatment in the early stages of disease has provided the impetus for generation of numerous gene expression datasets of intact tissue or RNA-seq of isolated cell populations and more recently the generation of multiple large scRNA-seq datasets comparing cellular phenotypes in inflamed and healthy intestinal mucosa ^7, 8, 17, 31-33^. The analysis of these datasets using dimensionality reduction algorithms such as UMAP produces a large array of putative subsets of both immune and non-immune cell types. However, there are two major issues that compromise interpretation. Firstly, the recovery of resident tissue macrophages by enzymatic disaggregation is substantially lower than their abundance *in situ* and is not necessarily representative. Based upon relative abundance of transcripts encoding known resident macrophage-specific proteins (CSF1R, C1Q), macrophages contribute around 10-15% of total human intestinal mRNA^34^. They are readily detected in total tissue mRNA. Secondly, the isolated macrophages that are profiled in scRNA-seq data are clearly activated during the process of tissue disaggregation as has been shown in isolated mouse macrophages from multiple tissues ^35^. Kong *et al*. ^8^ identified CCL3^+^CCL4^+^ macrophages as the most abundant population in both normal and inflamed mucosa, but neither *CCL3* nor *CCL4* mRNA is detectable in total RNA-seq data from normal ileum or colon in the FANTOM5 resource ^34^ nor in normal mucosa in our data, whereas *CSF1R* is readily detected. Similarly, Domanska *et al*. ^17^ and Elmentaite *et al*. ^32^ identified subsets of macrophages in normal human colon that expressed *IL1B*, numerous other pro-inflammatory chemokines and cytokines, and immediate early genes (e.g. *FOS, JUN*), none of which is detected by RNA-seq analysis of normal intestinal mucosa ^34^. Against this background, it is difficult to evaluate the predictive value of the GIMATS cellular module proposed by Martin *et al*. ^7^ as an indicator of pathogenesis.

As a consequence of random sampling each intestinal biopsy contains different proportions of structures and cell types enabling deconvolution. Network-based deconvolution of total RNA-seq data was used previously to extract signatures of immune cell types in solid tumors^23^. Similarly, the network analysis in **Table S4** highlights co-expression modules for various intestinal wall cell populations that can provide a framework for interpreting scRNA-seq data. **Cluster 20** includes markers of several distinct cell types in the intestinal crypts (*ASIC2, CHGA/B, DEFA/B, GCG, LGR5, REG3A, RFX6, SOX4*). Because crypts are well-defined structures, the individual cell types within them do not cluster separately. Conversely, monocyte and macrophage subpopulations vary in relative abundance in different locations within the intestinal wall. Two modules are consistent to some extent with resident lamina propria macrophages (**Cluster 21**) and submucosal macrophages (**Cluster 25**) identified by others ^17^ and with data from mouse ^35^. **Cluster 16** contains *CD14*, the key monocytic transcription factor, *CEBPB* and multiple monocyte-specific genes ^24^. **Cluster 8**, comprised of transcripts that share over-expression in all inflamed tissues, including draining lymph nodes, contains numerous inducible inflammatory genes associated with recruited monocytes.

The extensive published RNA-seq data on inflamed gut mucosa is limited in that it does not focus on AU, the earliest pathological lesion. **Table 1** highlights the distinct classes of transcripts that were increased in AU relative to PP or mucosa. The induction of immunoglobulin genes alongside an increase in B cell markers (e.g. *CD19, CD79, BTK*) indicates germinal center formation ^36^ and plasma cell differentiation to produce both IgA and IgG. This process may be driven by the up-regulation of key chemokines (*CCL19, CCL21, CXCL13*), their receptors (*CCR7, CXCR4*) and *TNFSF13B*, the transcript encoding B cell activating factor (BAFF).

Increased levels of transcripts for monocyte-specific genes (e.g. *CD14, NOD2, S100A8/9*; **Clusters 8** and **14**) indicates specific infiltration of these cells into the AU. The S100A8/9 complex, also known as calprotectin, is detectable in stools as well as serum of patients with IBD and has been used as a non-invasive biomarker of disease, likely associated with neutrophils in the intestinal lumen ^37^. Zollner *et al*. ^38^ showed that fecal calprotectin was well-correlated with another neutrophil-associated protein, lipocalin 2 (LCN2), that was also increased in AU (**Table 1**). Immunolocalisation revealed increases in both proteins in neutrophils and monocyte-macrophages in inflamed mucosa. *LCN2* was also induced in epithelial cells^38^ and in our RNA-seq data is around 5-10 fold more highly-expressed than S100A8/A9.

Monocyte recruitment may be linked to increased expression of multiple receptors implicated in trans-endothelial migration, including selectins (*SELE, SELP, SELL*), *PECAM1* and *ICAM1*^39^. Associated with this infiltration we see expression of transcripts encoding inducible cytokines such as IL1B, that are massively more inducible in monocytes than in monocyte-derived macrophages ^24^. This observation is consistent with our proposal that intestinal inflammation is triggered by hyper-responsiveness of monocytes to intestinal flora. The inducible response may also depend upon priming with interferon−γ (IFNG). **Cluster 27**, also over-expressed in AU, groups IFNG with multiple known IFNG target genes notably *CXCL9/10/11, IL12* and *STAT1* ^40^. Anti-IFNG antibody (fontolizumab) has been tested clinically in Crohn’s patients ^41^. Interestingly, *ACOD1* is induced in human monocytes and macrophages *in vitro* and functions to divert the citric acid cycle to produce itaconate ^24^. It has been highlighted as an IFNG target and potentially involved in IBD ^42^. However, *ACOD1* mRNA was barely detectable in AU or inflamed mucosa (max 18 TPM).

Our focus on AU also highlights the response of epithelial cells; transcripts associated with the stress response, **Cluster 4** and more specifically **Cluster 14**, were both strongly enriched in AU. A key observation herein is the profound increase in expression of CEACAM5/6/7 in AU (Table 2). These proteins are induced by bacterial toxins and provide the major portal for invasion by enteropathogenic *E*.*coli* ^43^. Other highly-induced genes are part of an antimicrobial defence also increased in IBD, including various mucins and the three trefoil factors, that contribute to non-specific barrier function. However, *MUC1*, the most highly-induced mucin in AU, was shown also to provide an inducible portal for bacterial uptake ^44^. Increased expression of *REG1A, LCN2, DUOX2* in inflamed mucosa was reported previously ^45^. DUOX2 generates hydrogen peroxide as an inducible defence against microorganisms. Interestingly, biallelic mutation of DUOX2 was recently described in a very early onset IBD patient ^46^. By contrast to rodents, in humans NOS2 (inducible nitric oxide synthase) is not expressed by macrophages ^47^. Conversely, *NOS2* mRNA and nitric oxide production is induced directly in human epithelial cells by invasive *E*.*coli, Salmonella* and *Shigella* species ^48^. Increased expression of IDO1 in epithelial cells has also been reported in ilea of Crohn’s disease patients ^49^.

Taken together, both the increased expression of inflammatory cytokines by recruited monocytes and associated alterations in epithelial gene expression in AUs strongly implicate a response to a microbial stimulus. In contrast, the absence of significant epithelial cell injury in the mucosa adjacent to AUs (Figure 2) argues against a global dysbiosis causing a breakdown in epithelial integrity as the initiating factor in the development of Crohn’s disease arising from AUs.

The analysis herein and previous studies ^16, 27^ provide very limited evidence for the selective enrichment or depletion of specific taxa in mucosal lesions including AU. This does not preclude a possible role for a microorganism as the initial trigger of focal inflammation in PP. Subsequent chronic inflammation may lead to disease-associated dysbiosis, but the evidence for this has been questioned in a recent systematic review of the published studies ^50^.

Our analysis of a relatively small dataset demonstrates that it is possible to extract information that complements, and in some cases clarifies, data generated by scRNA-seq. The results are consistent with the concept that CD pathology is triggered by a microbial stimulus at the site of lumenal antigen sampling. Genetic susceptibility may determine the nature of the host immune response to a specific “cognate” microbial trigger (which may differ between individuals) and/or regulate the nature, magnitude and duration of the response in those who progress to develop the full clinical manifestations of Crohn’s disease.

## Acknowledegements

This work was supported in part by an Australian National Health and Medical Research Council (NHMRC) Early Career Fellowship (APP1091221). DAH and KMSacknowledge core support from The Mater Foundation. The Translational Research Institute receives funding from the Australian Government. The authors thank the patients who participated in the study, and the staff at the Biological Research Facility, Australian National University, and the Canberra Hospital for their assistance. In addition, the authors would like to acknowledge Tom Freeman for bioinformatic support.

