## Supplementary material for "Transcriptomic signatures of host immune responses in aphthous ulcers, the earliest lesions of Crohn’s disease, suggest that a bacterial invasive challenge, rather than global dysbiosis, is the initiating factor": Table S1

| **Table S1**. Patient characteristics | | | | | |
| --- | --- | --- | --- | --- | --- |
|  | CD / Control | Tissue type | Sample size | Female / Male | Mean Age yrs (Std. dev) |
| Aphthous ulcer and Peyer’s patches samples and controls | CD | AU terminal ileum | 14 | 8F / 6M | 44.7 (7.4) |
|  | CD | Adjacent NM | 14 | 8F / 6M | 44.7 (7.4) |
|  | CD | AU large bowel | 7 | 6F / 1M | 44.7 (6.9) |
|  | Control | NM | 12 | 9F / 12M | 37.6 (12.7) |
|  | Control | PP | 8 | 5F / 3M | 39.1 (12.7) |
| Established inflammation | CD | Active | 21 | 9F / 12M | 44.7 (12.8) |
|  | CD | Inactive | 19 | 10F / 9M | 45.7 (15.0) |
|  | Control | NM | 13 | 7F / 6M | 47.7 (9.8) |
| Surgical resections | CD | Involved mucosa | 9 | 2F / 7M | 27.4 (14.5) |
|  | CD | Adjacent uninvolved mucosa | 7 | 2F / 5M | 26.7 (14.6) |
|  | CD | LN | 7 | 2F / 5M | 25.9 (10.6) |
|  | Control | NM | 9 | 5F / 4M | 57.0 (16.7) |
|  | Control | LN | 9 | 5F / 3M | 61.9 (11.7) |
