## Supplementary material for "Transcriptomic signatures of host immune responses in aphthous ulcers, the earliest lesions of Crohn’s disease, suggest that a bacterial invasive challenge, rather than global dysbiosis, is the initiating factor": Table S2

Supplementary Table 2. Quantitative real-time PCR Primers

|  | Forward 5’ to 3’ | Reverse 5’ to 3’ | R^2^ | Primer efficiency (%) |
| --- | --- | --- | --- | --- |
| GAPDH | CTTTTGCGTCGCCAG | TTGATGGCAACAATATCCAC | 0.99 | 111.9 |
| ACTB | GACGACATGGAGAAAATCTG | ATGATCTGGGTCATCTTCTC | 0.99 | 113.6 |
| TREM1 | ACAGATATCATCAGGGTTCC | CCTAGGGTACAAATGACCTC | 0.99 | 116.2 |
| FPR1 | ATCATTAGAGCCTGAGTCAC | TCTCCATCTTGTCTGCTC | 0.99 | 109.0 |
| OSMR | GTGTACAAGATTCTACTGGAC | GTTTCCCTTCCAAATAACAGG | 0.98 | 103.4 |
| TGM2 | CTTCATTTTGCTTCAACG | AGGATCCCATCTTCAAACTG | 0.99 | 107.8 |
| CEACAM6 | AACTCTTGGTATTACCCTCC | TGAGTTTTGTAATTCCAGCC | 0.99 | 109.0 |
| CEACAM4 | GCCTCACTTTTAACGTTCTG | TATAACCAGCAATGAGAGGG | 0.97 | 101.1 |
| S100A12 | GGGAATTGTCAATATCTTCCAC | CGACCTGTTCATCTTGATTAG | 0.99 | 106.8 |
| IGHA1 | GACACCTTCTCCTGCATGGT | ACATTGACATGGGTGGGTTT | 0.98 | 108.5 |
| TAP1 | CCTTCACTCGAAACTTAACTC | TGGTTCTGTTGGAAAAACTC | 0.99 | 103.0 |
| OSM | CCAGCTCCAGAAGCAGACAG | GCAGTGCTCTCTCAGTTTAGGAACAT | 0.98 | 101.4 |
| FCN1 | CATGCTTCAAACCTCAATGGTCT | GCCGCACTCCAGTTGATACC | 0.96 | 100.8 |
